# Mapping Amyloid Networks in Alzheimer’s Disease: A High-Order ICA Approach to Gray and White Matter Pathology

**DOI:** 10.1101/2025.03.21.644614

**Authors:** Nigar Khasayeva, Kyle M. Jensen, Cyrus Eierud, Helen Petropoulos, Allan I Levey, James Lah, Ruiyan Luo, Enrico Premi, Barbara Borroni, Jiayu Chen, Vince D. Calhoun, Armin Iraji

## Abstract

**INTRODUCTION:** Alzheimer’s Disease (AD) is a neurodegenerative disorder marked by gray matter (GM) changes driven by amyloid-beta (Aβ) plaques and neurofibrillary tangles. While GM alterations are well documented, spatially distinct patterns of homogeneous Aβ uptake and white matter (WM) involvement remain underexplored.

**METHODS:** We applied high-order independent component analysis (ICA) to 716 [18F]Florbetapir PET scans, identifying 80 GM and 13 WM networks. Diagnostic and cognitive associations were evaluated via statistical modeling.

**RESULTS:** Identified networks delineated a progression trajectory, with mild cognitive impairment (MCI) profiles in temporoparietal and frontal subdomains more closely aligned with AD than cognitively normal (CN) profiles. GM networks, including the hippocampal-entorhinal complex and precuneus, and WM networks, including the retrolenticular internal capsule, demonstrated robust associations with cognitive performance.

**DISCUSSION:** Our findings highlight the utility of high-order ICA in identifying reproducible Aβ networks and the contribution of WM networks, such as the posterior corpus callosum, in the early pathological landscape of AD.

## Introduction

Alzheimer’s disease (AD) is a preeminent cause of dementia, characterized by a gradual decline in cognitive functions such as memory, reasoning, and language, affecting approximately 32 million individuals worldwide [1], [2]. Given that the pathological processes underlying AD are believed to commence decades before clinical symptoms emerge [3], elucidating the neurobiological mechanisms during its prodromal stages is essential. Mild cognitive impairment (MCI) represents an intermediate phase between normal aging and dementia, with an annual conversion rate to AD dementia of 10–15% [4], [5].

AD pathology has been associated predominantly with gray matter (GM) abnormalities, particularly the aggregation of amyloid-beta (Aβ) plaques and neurofibrillary tangles, widely recognized as hallmark features of AD [6]. However, neuropathological and neuroimaging studies have shown that white matter (WM) degeneration may be an integral component of AD’s pathological cascade rather than a mere comorbidity [7], with changes such as axonal myelin degradation emerging early in the disease course, even before significant GM atrophy becomes evident [8]. This emphasizes the necessity of further investigating WM involvement in AD pathophysiology.

Positron Emission Tomography (PET) has transformed in vivo Aβ detection using amyloid-targeting agents such as [18F]Florbetapir ([18F]FBP), which permeate the blood-brain barrier and produce a radioactive signal that is detectable throughout the brain [9]. [18F]FBP is renowned for its high accuracy in detecting Aβ plaques [10]. It has been shown to detect uptake differences in cognitively normal (CN), MCI, and AD individuals [11], [12]. A recent study has revealed that [18F]FBP can also identify alterations in WM, particularly the generalized degradation of axonal myelin sheaths [7]. Myelin changes have been associated with cortical Aβ deposition in the preclinical stages of AD [8]. Despite these findings, WM alterations remain underexplored, and few studies have systematically examined the spatial organization of Aβ across the brain.

Conventional PET analysis methods typically rely on predefined regions of interest (ROI) to detect variations in tracer uptake [13]. These methods are labor-intensive, operator-dependent, and may miss heterogeneous uptake patterns that deviate from anatomical boundaries. ROI-based methods rely on a fixed number of predefined anatomical regions, which fail to encompass the entire brain space [14]. Voxel-based analysis [15] offers a more granular view of tracer distribution across the entire brain space. However, as a univariate approach, it analyzes each voxel in isolation, potentially overlooking inter-voxel relationships. It also remains susceptible to the low signal-to-noise ratio inherent to PET imaging, which can lead to misestimation of tracer uptake in voxels [16]. Both approaches may yield results that are unstable under multiple comparison correction. Multivariate data-driven methods like independent component analysis (ICA) address these challenges by integrating voxel-level information and capturing inter-subject covariance to reveal spatial patterns that represent regions with similar tracer uptake across subject groups [17].

ICA has shown promise in advancing PET-based research on AD. Prior studies have applied ICA to amyloid and tau PET data to identify network-level deposition [18], used [^18^F]Fluorodeoxyglucose ([^18^F]FDG) PET to identify age-related metabolic declines [19], and combined tracers using Parallel ICA to study language and visuospatial impairments [20]. Despite recent advances, PET studies have yet to fully exploit the potential of high-model-order ICA, which can isolate finer-grained, spatially distinct networks and enhance sensitivity to subtle disease-related changes [21].

In this study, we applied high-model-order ICA (model order = 100) to [18F]FBP PET data to extract both GM and WM networks and observe group-level differences in Aβ uptake between CN, MCI, and AD groups. We further assessed associations between these PET-derived networks and a range of clinical, cognitive, and neuropsychiatric measures to elucidate their relevance to AD symptomatology, which may also offer valuable insights into targeted treatment. This approach offers a comprehensive framework for characterizing the spatial architecture of Aβ deposition and its clinical implications in GM and particularly in WM, which remains an underexplored substrate of AD.

## Materials and Methods

### Participants

Data for this research were obtained from the ADNI database (adni.loni.usc.edu). ADNI is a longitudinal study designed to assess whether imaging modalities, such as PET and magnetic resonance imaging (MRI), combined with other biological markers, clinical evaluations, and neuropsychiatric assessments, can be integrated to measure the progression of MCI and early AD [22]. In this study, we focused on [18F]FBP PET imaging due to its established efficacy in capturing Aβ deposition.

The dataset comprised 716 subjects, each with two time points that included PET brain scans as well as corresponding demographic and clinical information. Subjects were classified into three groups based on their diagnostic status: CN individuals (N = 293), individuals with MCI (N = 274), and individuals with AD dementia (N = 149). For the primary analysis, one time point per subject was randomly selected, while the second time point was used to assess replicability. To account for potential confounding factors, we corrected the ICA loadings for site, age, and sex effects. Site was included as a covariate in the correction procedure to mitigate inter-site variability. Moreover, to ensure stable group estimates, only sites with at least 10 subjects were retained, resulting in a final sample of 624 subjects (Table 1). Age differences between groups were not significant, but given the established age-related nature of AD dementia, age was included in our correction procedure. Sex showed a significant difference between diagnostic groups and was therefore included as a covariate in our analysis.

**TABLE I.** Demographics of the study group after site correction.

| Diagnostic | N | Sex | N | Age | ADAS11 | MMSE | FAQ | CDRSB |
| --- | --- | --- | --- | --- | --- | --- | --- | --- |
|  |  |  |  | Mean (SD) |  |  |  |  |
| CN | 248 | Male | 104 | 76.63 (8.00) | 6.31 (3.38) | 28.76 (1.38) | 0.25 (0.95) | 0.04 (0.15) |
|  |  | Female | 144 | 73.81 (7.31) | 5.06 (2.62) | 29.12 (1.26) | 0.30 (0.83) | 0.10 (0.27) |
| MCI | 242 | Male | 142 | 76.49 (7.18) | 9.63 (4.78) | 27.73 (1.91) | 3.44 (4.55) | 1.58 (1.20) |
|  |  | Female | 100 | 74.68 (7.87) | 9.51 (4.46) | 27.96 (1.80) | 2.99 (3.65) | 1.54 (0.97) |
| AD<br>Dementia | 134 | Male | 70 | 77.11 (8.46) | 21.56 (8.45) | 22.06 (3.86) | 15.04 (7.71) | 5.25 (2.54) |
|  |  | Female | 64 | 74.07 (8.59) | 21.78 (8.60) | 21.92 (3.97) | 15.33 (7.18) | 5.23 (2.46) |

CN = Cognitively Normal; MCI = Mild Cognitive Impairment; AD = Alzheimer's Disease; ADAS-11 = Alzheimer's Disease Assessment Scale (11-item); MMSE = Mini-Mental State Examination; FAQ = Functional Activities Questionnaire; CDR-SB = Clinical Dementia Rating – Sum of Boxes.

### Image acquisition and preprocessing

All images were acquired using the [18F]FBP radiopharmaceutical and captured in four 5-minute frames, with slice thicknesses ranging from 1.0 mm to 3.4 mm. The four 5-minute PET frames were realigned utilizing PETPrep_HMC (github.com/mnoergaard/petprep_hmc) and registered to the T1-weighted MRI scans. Afterward, the four realigned and registered frames were spatially normalized to the MNI305 template [23] and averaged to produce a single motion-corrected PET image for each participant. Then, PET voxel intensities were normalized to standardized uptake value ratios (SUVR) using the cerebellar cortex as the reference region [24]. The cerebellar cortex is frequently employed as a reference region due to the binding of [18F]FBP PET to Aβ plaques, which are notably less prominent in the cerebellum [25]. Finally, the images were smoothed using a 10 mm Gaussian kernel [26].

### [18F]FBP PET networks identification with high-model-order independent component analysis

To identify spatially distinct patterns of amyloid-β uptake, we applied ICA, a blind-source separation method [27] commonly utilized to identify maximally spatially independent patterns that covary across subjects or time. Specifically, we used high-model-order ICA with the number of components set to 100. Prior to decomposition, subject-level PET data were concatenated and reduced using principal component analysis (PCA), followed by ICA using the Infomax algorithm [28]. To obtain reliable independent components (ICs), we utilized the ICASSO module of Group ICA of fMRI Toolbox [29] (GIFT v4.0.5.27, github.com/trendscenter/gift), which runs ICA multiple times using bootstrapping and random initialization. ICASSO uses clustering to identify the most stable components as centroids, ensuring reproducibility across iterations [30]. Finally, ICA outputs the spatial components and their corresponding loadings for subjects. In our analysis, we specifically focused on these loadings derived from selected components, i.e., networks, examining how variations in these network loadings relate to Aβ accumulation and different stages of cognitive impairment.

Of the 100 components identified, many corresponded to GM and WM networks, while others reflected ventricle-related or artifactual components. We then proceeded with the GM and WM networks, manually labeling them based on their spatial distribution. GM labels followed the NeuroMark 2.2 naming convention [31], a data-driven atlas of canonical brain networks derived from over 100,000 resting-state fMRI datasets [32], preserving its domain and subdomain structure. WM labels were assigned using the JHU white matter atlas as the anatomical reference [33], which served as an anatomical reference for identifying the spatial distribution of WM networks. Manual labeling was performed by two raters, authors NK and KJ, and validated by the senior author AI.

### Statistical analysis for cross-group comparison and relationship between PET network loadings and cognitive/neuropsychiatric symptoms

To minimize the influence of nuisance variables and ensure biologically meaningful findings, we regressed out site, age, and sex from the PET network loadings. For the group-level comparisons, the adjusted loadings were then entered into two-sample t-tests to evaluate differences between CN, MCI, and AD. Multiple comparisons correction employed the Benjamini–Hochberg false discovery rate (FDR) [34], with a significance threshold of α = 0.05. The same analyses were repeated in the replicate set.

To examine the translational value of our findings, we tested associations between adjusted network loadings and cognitive/neuropsychiatric measures from the ADNI database, restricting analyses to networks that showed significant diagnostic differences. Partial correlation was applied to both loadings and behavioral scores, with the diagnostic group included as a covariate. This step was essential to prevent Simpson’s paradox [35], where group differences could otherwise obscure or invert brain–behavior relationships. Analyses were performed in the combined MCI and AD group (MCI+AD), focusing on four key clinical measures: Alzheimer’s Disease Assessment Scale 11-item cognitive subscale (ADAS11) [36] for cognitive evaluation (scores ranging from 0 for no impairment to 70 for maximum impairment), the Clinical Dementia Rating Sum of Boxes (CDRSB) [37] for global assessment of dementia severity (ranging from 0 for no impairment to 18 for severe impairment), the Functional Activities Questionnaire (FAQ) [38] for evaluating daily living activities (ranging from 0 for independence to 30 for maximum dependence), the Mini-Mental State Examination (MMSE) [39] for cognitive assessment (with 30 indicating no impairment and 0 maximum impairment). The same analyses were also repeated in the replicate set.

## Results

### Estimated networks across the Alzheimer’s disease course

#### Gray matter networks

Our analysis identified 80 PET-derived gray matter (GM) networks, which showed spatial correspondence with established GM functional networks [31], [32]. These networks were organized into seven major domains and further subdivided into 14 subdomains, including cerebellar (CB), visual-occipitotemporal (VI-OT), visual-occipital (VI-OC), paralimbic (PL), subcortical-extended hippocampal (SC-EH), subcortical-extended thalamic (SC-ET), subcortical-basal ganglia (SC-BG), sensorimotor (SM), higher cognition-insular-temporal (HC-IT), higher cognition-temporoparietal (HC-TP), higher cognition-frontal (HC-FR), and the triple network domain: central executive (TN-CE), default mode (TN-DM), and salience (TN-SA) (Figure 1). Comparison with the fMRI NeuroMark 2.2 template showed high spatial similarity (Pearson’s r > 0.6) in the CB, VI-OT, SC-BG, and TN-SA, while other domains exhibited lower similarity, reflecting more distinct characteristics between [18F]FBP PET and fMRI data.

**Figure 1.**
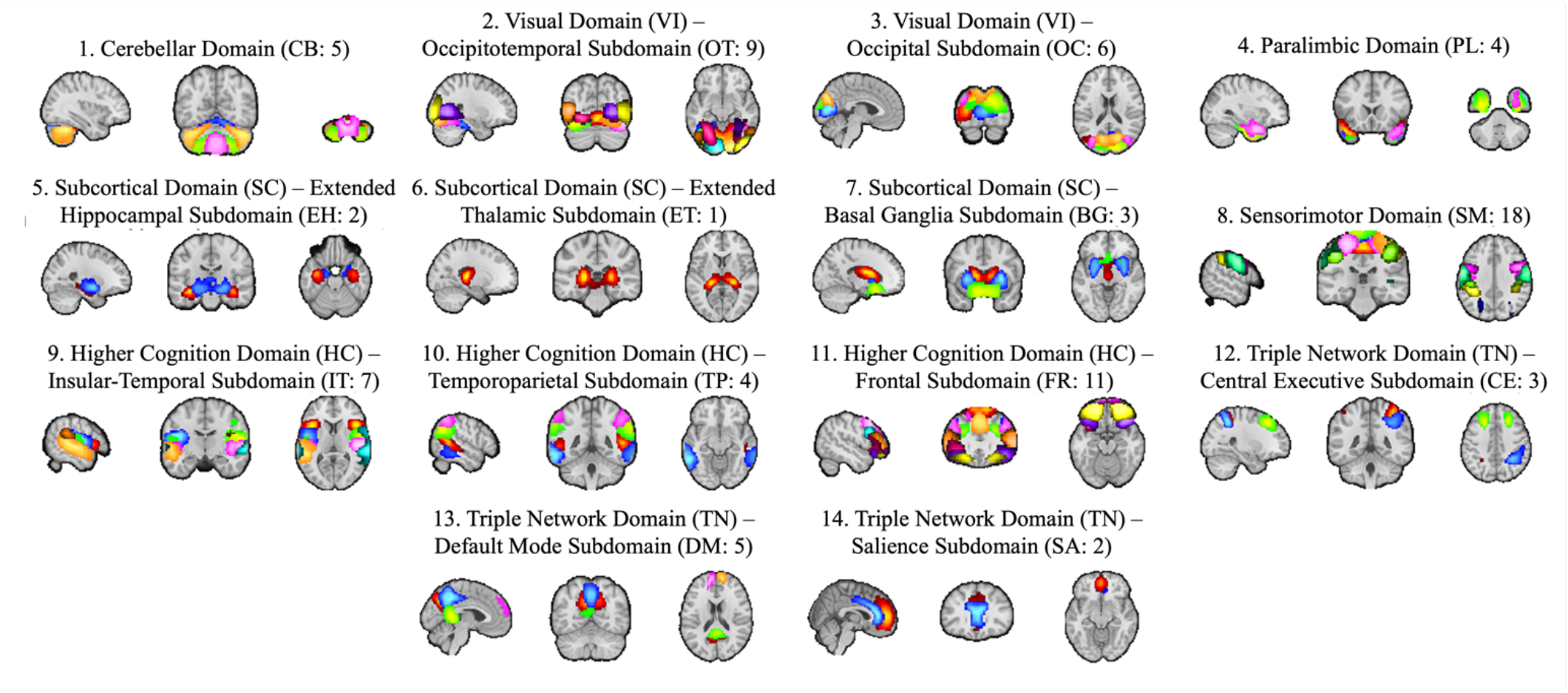
Spatial maps of 80 PET gray matter networks extracted from [18F]Florbetapir PET using independent component analysis, categorized into seven domains and 14 subdomains: 1) Cerebellar, 2) Visual-Occipitotemporal, 3) Visual-Occipital, 4) Paralimbic, 5) Subcortical-Extended Hippocampal, 6) Subcortical-Extended Thalamic, 7) Subcortical-Basal Ganglia, 8) Sensorimotor, 9) Higher Cognition-Insular-Temporal, 10) Higher Cognition-Temporoparietal, 11) Higher Cognition-Frontal, 12) Triple Network-Central Executive, 13) Triple Network-Default Mode, and 14) Triple Network-Salience.

#### White matter networks

We also identified 13 PET WM networks, each showing spatial correspondence to known WM tracts. These included the retrolenticular internal capsule (RICap), anterior corpus callosum (ACoCa), anterior corona radiata (ACRad), posterior corpus callosum (PCoCa), superior longitudinal fasciculus (SLFas), posterior corona radiata (PCRad), posterior cingulum (PCing), tapetum (Taptm), posterior thalamic radiation (PTRad), anterior middle cerebellar peduncle (AMCbP), cerebral peduncle (CePed), corticospinal tract (CSpTr), and posterior middle cerebellar peduncle (PMCbP) (Figure 2).

**Figure 2.**
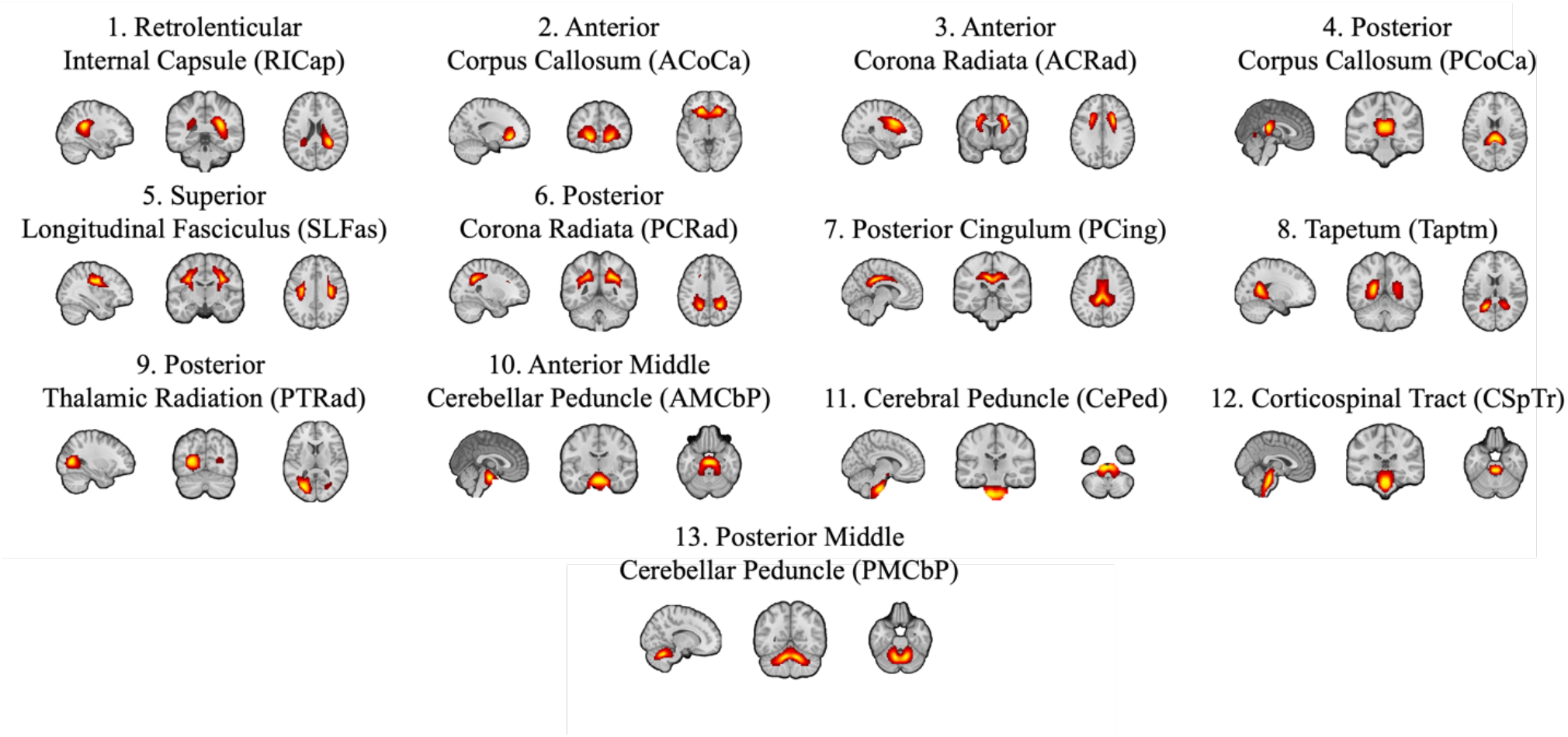
Spatial maps of 13 PET white matter networks extracted from [18F]Florbetapir PET using independent component analysis, including 1) Retrolenticular Internal Capsule, 2) Anterior Corpus Callosum, 3) Anterior Corona Radiata, 4) Posterior Corpus Callosum, 5) Superior Longitudinal Fasciculus, 6) Posterior Corona Radiata, 7) Posterior Cingulum, 8) Tapetum, 9) Posterior Thalamic Radiation, 10) Anterior Middle Cerebellar Peduncle, 11) Cerebral Peduncle, 12) Corticospinal Tract, and 13) Posterior Middle Cerebellar Peduncle.

All identified GM and WM networks exhibited high stability, with ICASSO cluster quality index (Iq) exceeding 0.8.

### Cross-group comparison of network loadings

#### Gray matter networks

The analysis of [18F]FBP PET data included comparisons across seven domains and 14 subdomains, with significant differences identified in numerous GM PET networks. These network-level findings reflected systematic progression in amyloid burden, with MCI consistently emerging as a transitional state between CN and AD. Trajectory analysis was operationalized by comparing the absolute differences between group means of adjusted network loadings for adjacent diagnostic pairs. By this criterion, HC-TP and HC-FR most consistently placed MCI closer to AD than to CN, with the same pattern observed intermittently in VI-OT, VI-OC, SC-ET, TN-CE, and TN-DM. In contrast, other subdomains more often placed MCI closer to CN than to AD.

Within the diagnostic trajectory from CN to MCI to AD dementia, the contrast between AD and CN showed the largest difference, while the AD vs. MCI and MCI vs. CN comparisons demonstrated fewer, though still notable, distinctions. To evaluate the robustness of these effects, we additionally tested all findings in an independent replicate set comprising randomly selected second scans from the same subjects. Specifically, in Figure 3a-c, 67 networks exhibited significant differences between AD dementia and CN, 58 networks between AD dementia and MCI, and 46 networks between MCI and CN. To evaluate the robustness of these effects, we additionally tested all findings in an independent replicate set. Of these, 58, 50, and 25 networks, respectively, showed significant effects that persisted in the replicate set.

**Figure 3.**
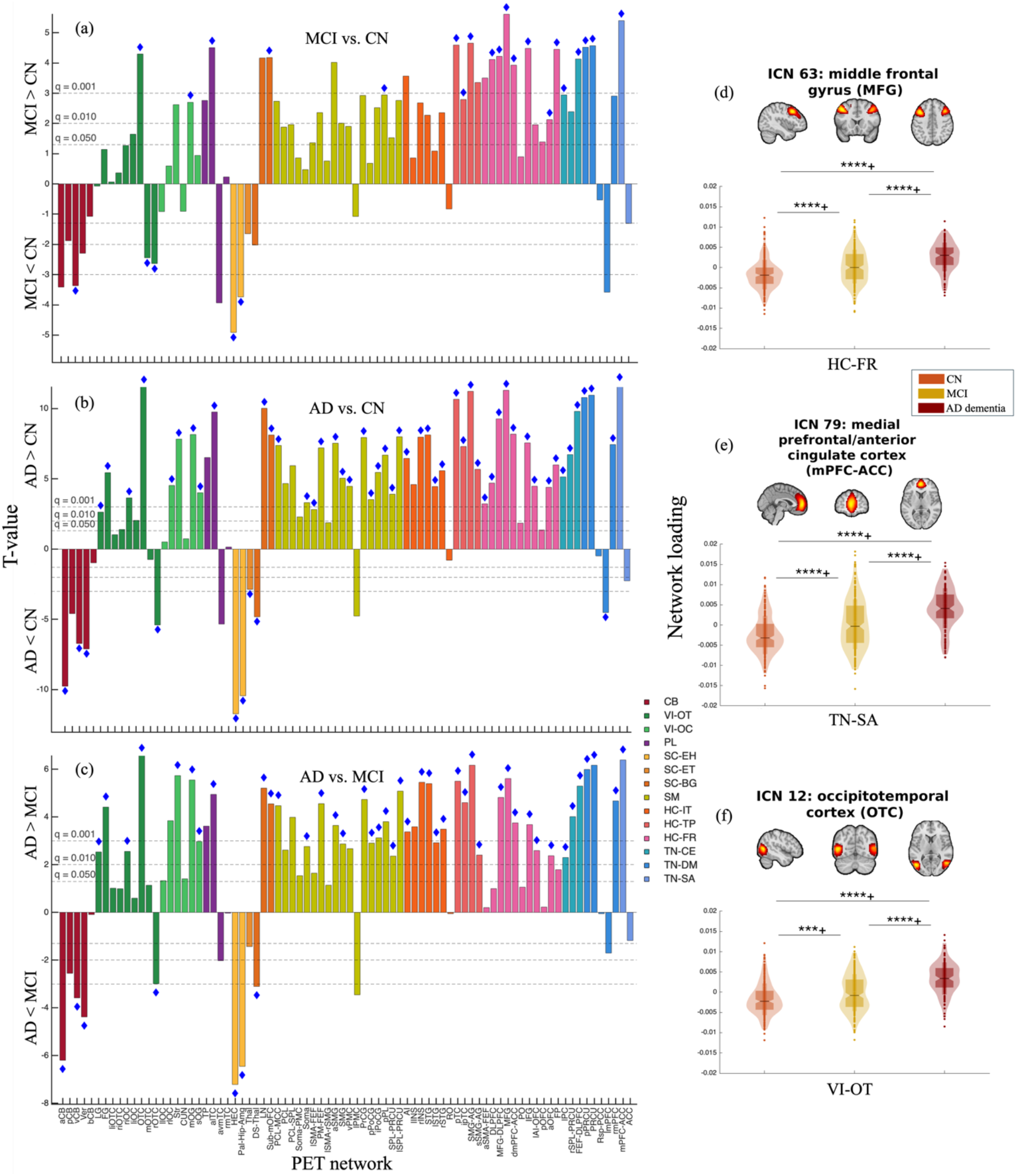
(a, b, c) Bar plots depicting t-values from two-sample t-tests on adjusted [18F]Florbetapir PET gray matter network loadings, extracted through high-order independent component analysis. The comparisons shown are Mild Cognitive Impairment (MCI) vs. Cognitively Normal (CN), Alzheimer’s Disease (AD) vs. CN, and AD vs. MCI, respectively, where positive t-values indicate that the first group in the comparison has higher network loadings, and negative t-values indicate the second group has higher loadings. (d, e, f) Violin plots illustrating network loading differences among CN, MCI, and AD groups for the most significantly affected subdomains in each pairwise comparison: HC-FR (Higher Cognition–Frontal), TN-SA (Triple Network– Salience), and VI-OT (Visual–Occipitotemporal), respectively. ****q < 0.0001, ***q < 0.001, **q < 0.01, *q < 0.05, corrected for FDR; ◆ indicates a significant effect in the replicate set.

We next examined the most affected subdomains across diagnostic comparisons. For the MCI vs. CN comparison, the most affected subdomains, with the strongest effect observed in the HC-FR (Middle Frontal Gyrus, Figure 3d), were followed by TN-SA, SC-EH, HC-TP, and TN-DM. These subdomains were significant in the discovery set and replicable in the replicate set. For the AD vs. CN comparison, the most affected subdomains, led by the TN-SA (Medial Prefrontal/Anterior Cingulate Cortex, Figure 3e), were followed by the VI-OT, SC-EH, HC-FR, HC-TP, TN-DM, and SC-BG. These subdomains were significant in the discovery set and in the replicate set. For the AD vs. MCI comparison, the most affected subdomains, with the greatest effect seen in the VI-OT (Occipitotemporal Cortex, Figure 3f), were followed by the SC-EH, TN-SA, CB, HC-TP, and TN-DM. These subdomains were significant in both the discovery and replicate sets.

To complement these subdomain-level findings, we also studied broader network-level effects. Within the CB, the anterior cerebellum and vermis showed highly significant differences in AD vs. CN (q < 0.0001) and AD vs. MCI (q < 0.0001), indicating that these networks become affected primarily at the AD stage, consistent with later involvement. Within the VI-OT, the fusiform gyrus network demonstrated highly significant differences in both AD vs. CN (q < 0.0001) and AD vs. MCI (q < 0.0001) comparisons, but no significant differences were found in the MCI vs. CN comparison (q > 0.05), suggesting later involvement. Within the VI-OC, the striate cortex and middle occipital gyrus displayed increasing significance throughout all comparisons as the disease progressed, with highly significant differences in AD vs. CN (q < 0.0001) and AD vs. MCI (q < 0.0001), consistent with progressive engagement as disease severity increases. Within the PL, the anterior lateral temporal cortex exhibited high significance (q < 0.0001) across all stages, underscoring its critical role in the progression from CN to MCI to AD dementia.

In the SC-EH, the hippocampal–entorhinal complex showed highly significant differences in all comparisons: MCI vs. CN (q < 0.0001), AD vs. CN (q < 0.0001), and AD vs. MCI (q < 0.0001). The pallidum/hippocampus/amygdala also showed strong effects in MCI vs. CN (q < 0.001), AD vs. CN (q < 0.0001), and AD vs. MCI (q < 0.0001). These findings underscore their major involvement throughout the disease progression. In the SC-ET, the thalamus did not show significant differences in the AD vs. MCI and MCI vs. CN comparisons (q > 0.05). However, significant differences were observed in the AD vs. CN comparison (q < 0.01), suggesting lower tracer accumulation, becoming detectable only when contrasting the two extremes of the disease trajectory.

Within the SC-BG, the lentiform nucleus network showed significant differences across all comparisons (q < 0.001). Its consistent involvement in the MCI vs. CN, AD vs. MCI, and AD vs. CN comparisons highlights the lentiform nucleus network as a key player throughout disease progression.

In the SM, the most prominent alterations were observed in the paracentral lobule/middle cingulate cortex, precentral gyrus, posterior parietal lobe, and premotor cortex, all showing robust significance in AD vs. CN (q < 0.0001) and AD vs. MCI (q < 0.001). The precentral gyrus and posterior parietal lobe demonstrated strong effects across all comparisons, including MCI vs. CN, underscoring their progressive involvement. In contrast, the somatosensory cortex, superior parietal lobe, and ventral/lateral premotor cortices showed no early differences in MCI vs. CN (q > 0.05) but became highly significant in AD vs. CN and AD vs. MCI, consistent with later involvement. Notably, the anterior supramarginal gyrus exhibited significance even in MCI vs. CN (q < 0.001), suggesting a potential role in early dysfunction.

In the HC-IT, the anterior insula and right insula emerged as key regions showing progressive disruption, with strong significance across all comparisons. The left insula showed no early-stage differences in MCI vs. CN (q > 0.05) but was highly significant in AD vs. CN (q < 0.0001) and AD vs. MCI (q < 0.001), suggesting later-stage involvement. The superior temporal gyrus, particularly in the right hemisphere, also demonstrated progressive decline with significance across all comparisons (q < 0.05), whereas its left counterpart showed alterations only at more advanced stages. Within the HC-TP, the posterior temporal cortex network and supramarginal/angular gyrus networks demonstrated highly significant differences in all comparisons (q < 0.0001), indicating their critical role in gradual disease progression. Within the HC-FR, the middle frontal gyrus/dorsolateral prefrontal cortex, and inferior frontal gyrus across all stages (q < 0.001), indicating progressive involvement.

Within the TN-CE, the frontal eye field/dorsolateral prefrontal cortex demonstrated high significance across all stages (q < 0.0001), reflecting progressive involvement. Within the TN-DM, the posterior precuneus and precuneus showed highly significant differences across all comparisons (MCI vs. CN, AD vs. CN, and AD vs. MCI; q < 0.0001), consistent with early and ongoing involvement. The left medial prefrontal cortex was significant in MCI vs. CN (q < 0.001) and highly significant in AD vs. CN (q < 0.0001) but not in AD vs. MCI (q > 0.05), suggesting early involvement with limited additional progression. Within the TN-SA, the medial prefrontal/anterior cingulate cortex was highly significant across all comparisons (q < 0.0001), indicating progressive involvement in disease stages.

The overall trends exhibited a gradient of changes in GM network loadings from CN to MCI to AD, with specific subdomains and their associated networks showing distinct progression patterns. The CB, SC-EH, and SC-ET exhibited predominantly decreased loadings as the disease progressed from CN to MCI to AD. In contrast, other subdomains displayed predominantly increasing patterns with advancing disease stages.

#### White matter networks

Expanding the analysis to WM PET networks, shifts in WM network loadings across the CN, MCI, and AD dementia groups revealed a comparable trajectory, highlighting MCI as a critical intermediate pattern in the gradual decline of white matter integrity. Notably, in ACoCa and PTRad, MCI values were closer to AD than to CN, suggesting greater alteration in these networks at the MCI.

As shown in Figure 4, we identified 11 WM networks that differentiated AD from CN, 10 that distinguished AD from MCI, and eight that separated MCI from CN. Importantly, seven networks (RICap, ACRad, PCoCa, SLFas, PCRad, CSpTr, and PMCbP) consistently showed significant differences across all comparisons. Overall, seven networks consistently demonstrated differences across all comparisons. Most of these effects were replicated in the independent dataset, further supporting the robustness and reproducibility of WM networks in capturing disease-specific patterns across diagnostic groups.

**Figure 4.**
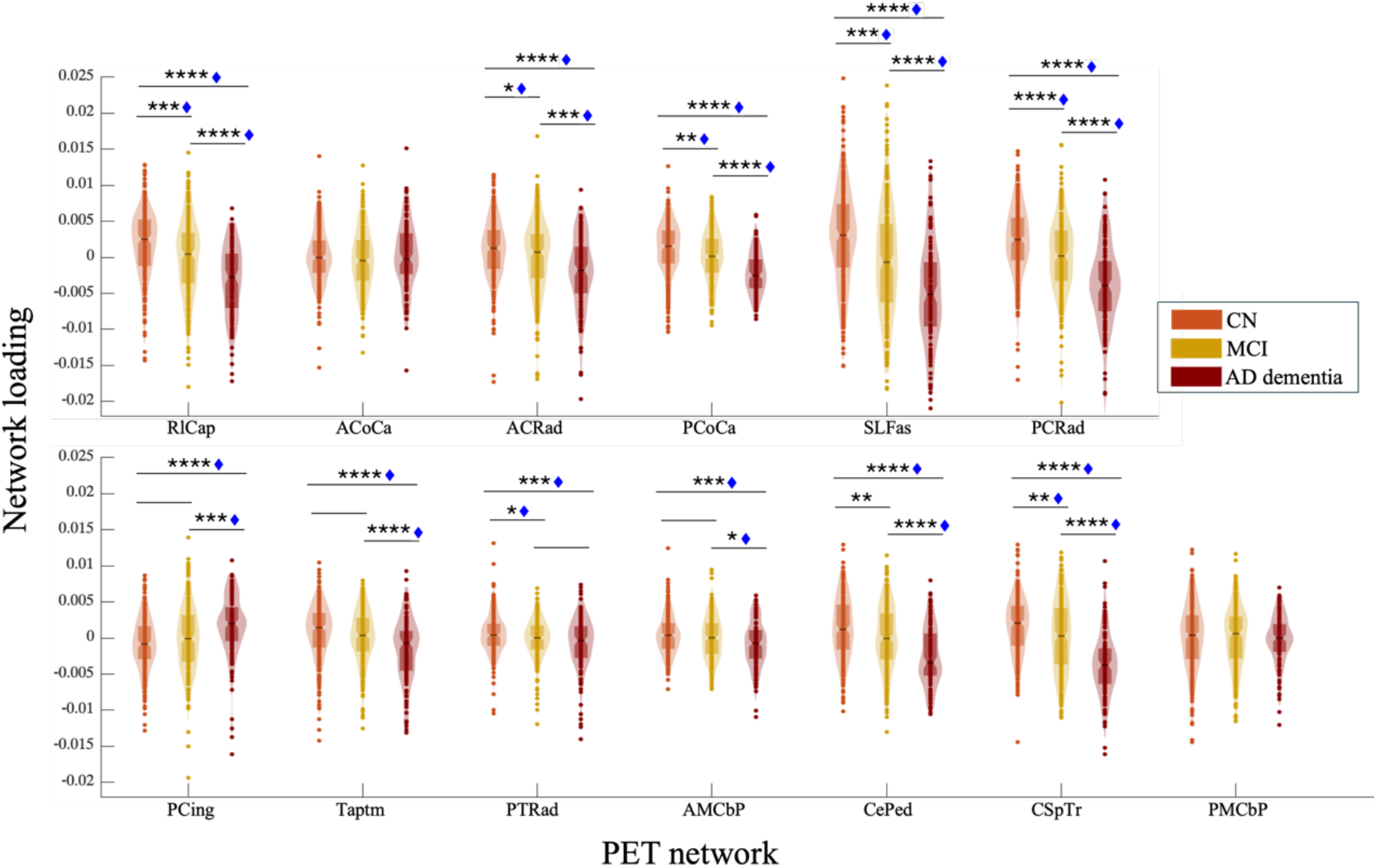
Violin plots showing PET white matter network loading differences between the three diagnostic groups: cognitively normal (CN), mild cognitive impairment (MCI), and Alzheimer’s disease (AD) dementia for Retrolenticular Internal Capsule (RICap), Anterior Corpus Callosum (ACoCa), Anterior Corona Radiata (ACRad), Posterior Corpus Callosum (PCoCa), Superior Longitudinal Fasciculus (SLFas), Posterior Corona Radiata (PCRad), Posterior Cingulum (PCing), Tapetum (Taptm), Posterior Thalamic Radiation (PTRad), Anterior Middle Cerebellar Peduncle (AMCbP), Cerebral Peduncle (CePed), Corticospinal Tract (CSpTr), and Posterior Middle Cerebellar Peduncle (PMCbP). ****q < 0.0001, ***q < 0.001, **q < 0.01, *q < 0.05, corrected for FDR; ◆ indicates a significant effect in the replicate set.

### Association with cognitive and neuropsychiatric variables

#### Gray matter networks

A partial correlation analysis was conducted for the MCI+AD group on PET-derived GM networks that exhibited significant diagnostic differences (Figure 5a). ADAS11 scores exhibited the most widespread correlations, spanning all subdomains except SC-ET and SC-BG. MMSE scores demonstrated significant associations with the VI-OT, SC-EH, SM, and TN-DM. FAQ scores correlated with the SC-EH and VI-OC, while CDRSB scores exhibited significant correlations with the SC-EH.

**Figure 5.**
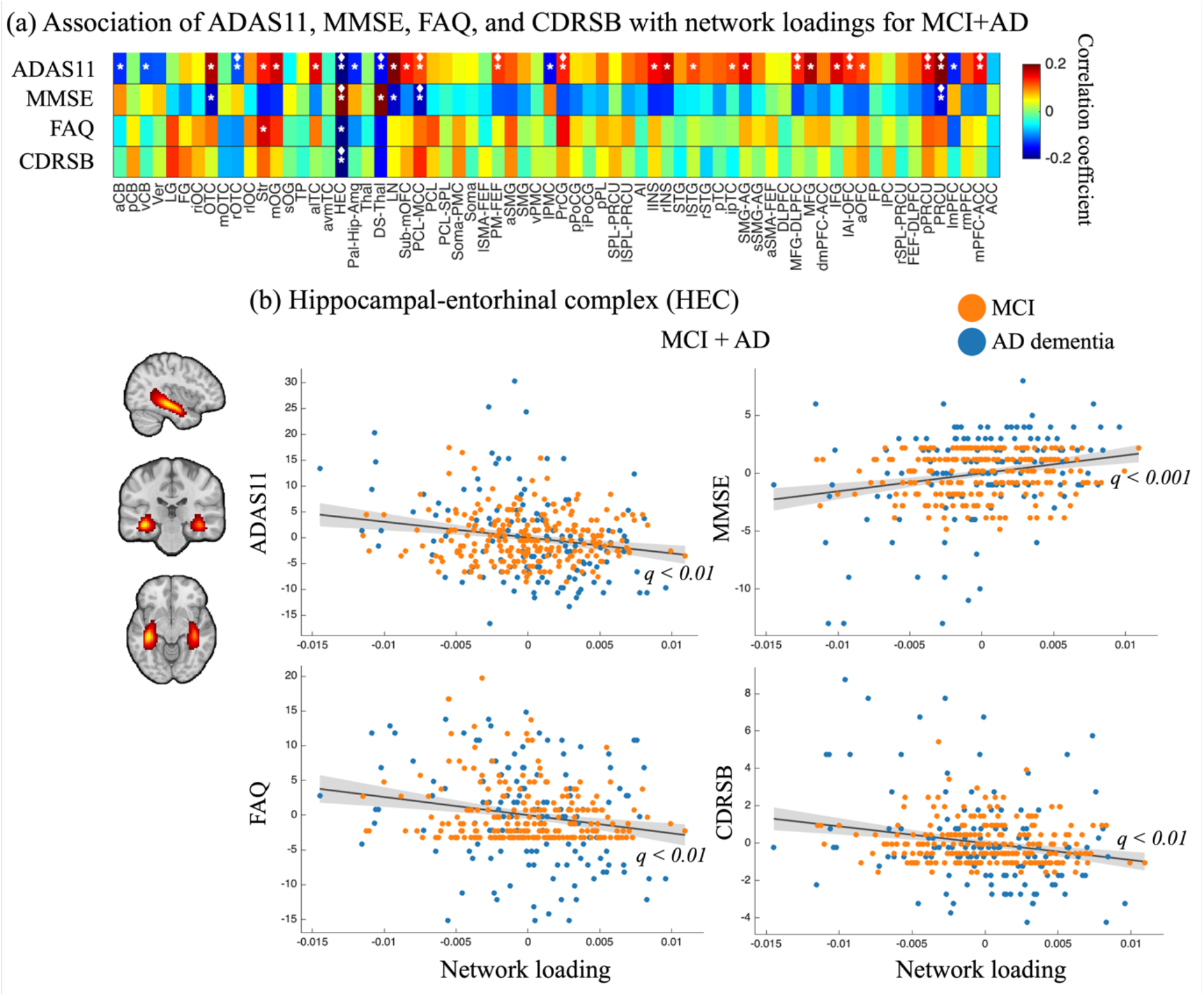
Partial correlation analysis between standardized PET gray matter network loadings and clinical measures—Alzheimer’s Disease Assessment Scale-Cognitive Subscale (ADAS11), Mini-Mental State Examination (MMSE), Functional Activities Questionnaire (FAQ), and Clinical Dementia Rating Sum of Boxes (CDRSB) for the combined group of mild cognitive impairment (MCI) and Alzheimer’s disease (AD) dementia. The analysis was performed across all 80 extracted networks. Significant correlations (q < 0.05), corrected for false discovery rate (FDR), are marked with star symbols; ◆ indicates a significant effect in the replicate set. (a) Associations between ADAS11, MMSE, FAQ, CDRSB, and network loadings across networks that showed significant group effect, (b) The fitted associations for extended hippocampal subdomain-hippocampal–entorhinal complex (HEC), demonstrating the relationship between network loadings and cognitive scores.

Among these, the SC-EH, specifically the hippocampal–entorhinal complex, exhibited consistent significance across all four measures, with ADAS11, MMSE, and CDRSB associations (Figure 5b) also replicated in the independent set. Additionally, within the TN-DM, the precuneus, and within the SM, the paracentral lobule/middle cingulate cortex, exhibited replicated significance for ADAS11 and MMSE.

#### White matter networks

A partial correlation analysis was also conducted for the MCI+AD dementia group on PET-derived WM networks that exhibited significant diagnostic differences (Figure 6a). ADAS11 scores exhibited the most widespread correlations, spanning all subdomains except PCing and AMCbP. MMSE scores demonstrated significant associations with the RICap, PCoCa, PCRad, Taptm, and CSpTr. FAQ scores were correlated with the RICap. ACRad, PCRad, PTRad, and CSpTr. CDRSB scores exhibited significant correlations with the RICap and the PCRad.

**Figure 6.**
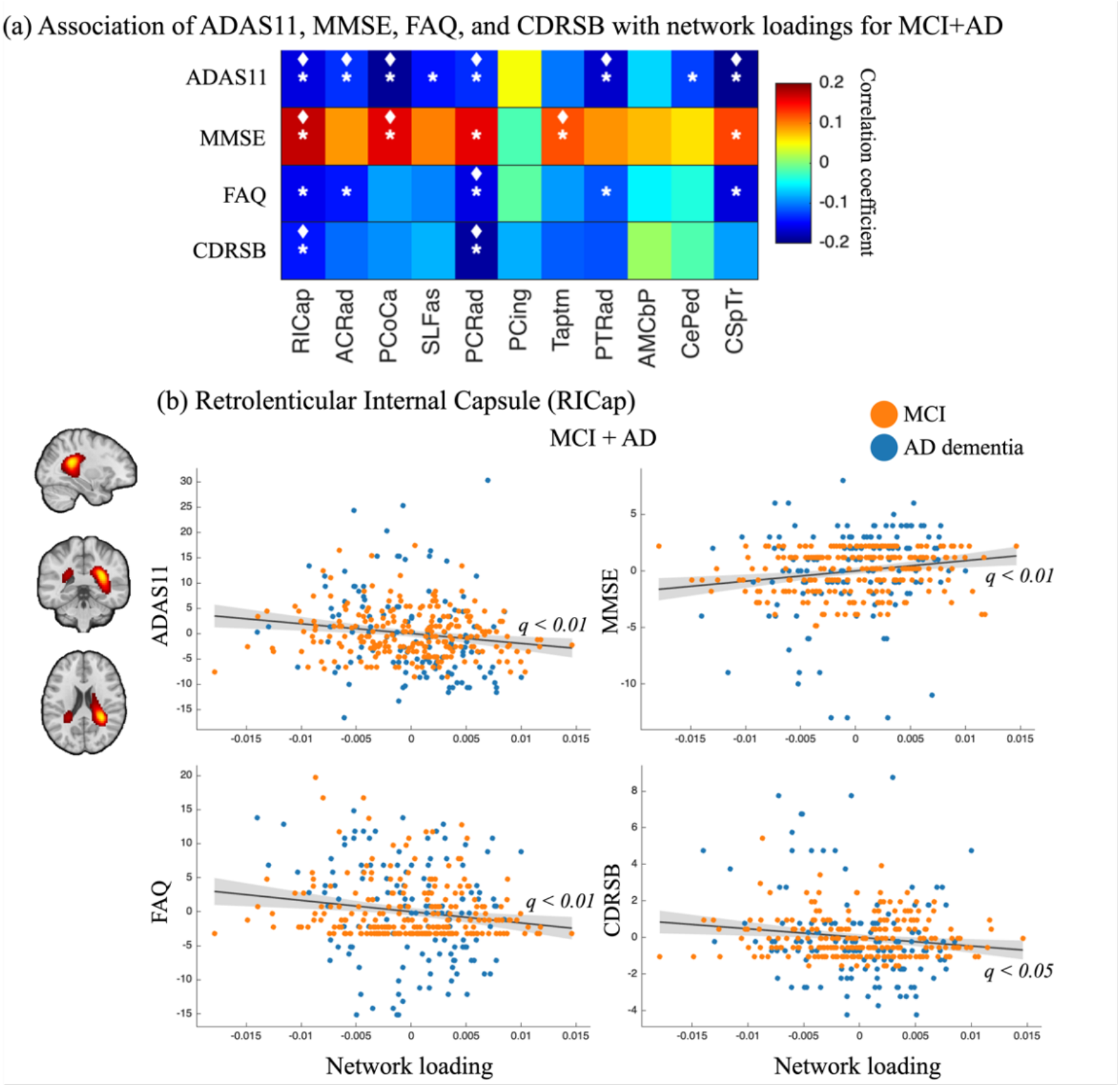
Partial correlation analysis between standardized PET white matter network loadings and clinical measures— Alzheimer’s Disease Assessment Scale-Cognitive Subscale (ADAS11), Mini-Mental State Examination (MMSE), Functional Activities Questionnaire (FAQ), and Clinical Dementia Rating Sum of Boxes (CDRSB) for the combined group of mild cognitive impairment (MCI) and Alzheimer’s disease (AD) dementia. The analysis was performed across all 13 extracted networks. Significant correlations (q < 0.05), corrected for false discovery rate (FDR), are marked with star symbols; ◆ indicates a significant effect in the replicate set. (a) Associations between ADAS11, MMSE, FAQ, CDRSB, and network loadings across all networks, (b) The fitted association for the Subcortical subdomain—retrolenticular internal capsule (RICap), demonstrating the relationship between network loading and cognitive scores.

Among these, the RICap (Figure 6b) exhibited consistent significance across all four measures, with ADAS11, MMSE, and CDRSB associations also replicated in the independent set. The PCRad also demonstrated consistent associations with all four measures, and ADAS11, FAQ, and CDRSB associations were replicated in the independent set.

## Discussion

We applied high-model-order ICA to investigate the neurobiological mechanisms underlying AD through the analysis of [18F]FBP PET data. By leveraging ICA, we identified spatially distinct brain networks that may offer novel insights into the progression of AD, particularly concerning amyloid deposition patterns and their effects on cognitive function. In earlier work [40], we used lower-model-order ICA on the same dataset to examine group differences between CN and AD, revealing effects in the salience, default mode, and temporal networks. In a separate study [41], we examined WM covariance patterns, showing that WM integrity provides meaningful diagnostic disparities across CN, MCI, and AD dementia. Building on these foundations, the use of high-model-order ICA not only recapitulates these known findings but also reveals more fine-grained and previously unrecognized network alterations that may elude lower-order methods. Our findings emphasize the role of both GM and WM alterations in AD pathology, with significant changes observed across multiple brain regions.

The observed similarity between PET and fMRI networks highlights the preservation of coherence across imaging modalities despite fundamental differences in their underlying physiological signals. While early-phase PET is often used as a proxy for cerebral perfusion and neurodegeneration (akin to FDG-PET), high-order ICA can extract comparable network-level information even from traditional amyloid PET imaging. Interestingly, dual-phase PET using both amyloid and tau tracers [42], [43] has demonstrated that the early phase reliably captures brain patterns closely aligned with FDG-PET, serving as a marker of cerebral perfusion and neurodegeneration. Variability in network correspondence may stem from methodological distinctions, differences in metabolic and functional architecture, or sample characteristics. Nevertheless, the ability of PET networks to recapitulate the established domain and subdomain structure of NeuroMark 2.2 underscores the robustness of [18F]FBP PET ICA in delineating meaningful neurobiological networks relevant to AD research.

In CB, highly significant differences were observed in the anterior cerebellum and vermis in AD compared to CN and MCI groups, suggesting late-stage amyloid accumulation in these regions. While often considered peripheral in cognitive research, the cerebellum plays a crucial role in motor coordination, balance, and mental processes such as working memory and attention. The observed pattern of late-stage amyloid accumulation suggests a potential role for the cerebellum in the widespread cognitive and motor decline that characterizes advanced AD [44], contributing to symptoms like gait disturbances and impaired coordination.

In the VI-OT, the fusiform gyrus is a specialized region for visual processing, including facial recognition and object identification [45]. The lack of an early effect suggests that the amount of amyloid accumulation or its detrimental effect on network function in the fusiform gyrus doesn’t reach a detectable threshold until the disease has advanced to the AD stage. This aligns with clinical observations that visual processing impairments, such as prosopagnosia (inability to recognize faces) or visuospatial deficits, are more prominent features of moderate to severe AD, rather than its earliest, most subtle stages. In the VI-OC, we found significant differences in the striate cortex and middle occipital gyrus across all comparisons, indicating both early involvement and progressive disruption of visual processing regions as the disease advances. In previous studies, considerable Aβ deposition was observed in the occipital lobes in patients with AD [46], [47], [48]. Given that individuals with occipital amyloid accumulation exhibit visuospatial and visual memory impairments even in the MCI stage, our findings reinforce the idea that occipital Aβ accumulation can be an overlooked marker of an earlier-onset or more aggressive disease course [49].

In the SC-ET, the thalamic alterations were relatively subtle, with lower tracer accumulation. Whole-thalamus analyses can mask the vulnerability of specific nuclei, since early pathology primarily affects limbic and associative nuclei while motor and sensory nuclei remain relatively preserved. This dilution of the true extent of degeneration may explain the lack of significance in early comparisons, with statistical differences emerging only when contrasting AD with CN, where neurodegeneration is more widespread. Given the thalamus’s role as a relay center crucial for sustaining attention and consciousness [50], even subtle changes in its associative nuclei could have important implications for cognitive decline in AD. The SC-BG is involved in motor control, executive decision-making, and reward or aversion-emotional stimulation [51]. In pathologic studies, the presence of amyloid plaques in the striatum, the largest structure of the basal ganglia, predicts a prevalence of dementia and clinicopathological AD [52]. The presence of striatal plaques was also correlated with lower scores on several neuropsychological tests assessing memory [53].

In the HC-FR, significant differences across all comparisons indicate that these frontal regions as the middle frontal gyrus/dorsolateral prefrontal cortex and inferior frontal gyrus, which are essential for higher executive functions like planning, decision-making, and cognitive flexibility [54], are under continuous pathological effect throughout the disease course. The most pronounced differences, as seen in the AD vs. CN comparison, align with neuropathological staging models that identify the frontal cortex as one of the most susceptible regions for Aβ deposition in AD [55], [56], [57]. This progressive compromise of the frontal regions directly contributes to the executive dysfunction and impaired cognitive control that are hallmark symptoms of AD.

Within the TN, the posterior precuneus and precuneus exhibited significant differences in AD dementia vs. CN, consistent with research documenting Aβ accumulation and metabolic decline in the default mode network in AD. The default mode is involved in introspective and self-referential processes and has been consistently implicated as a network affected by AD [58]. It has been suggested that Aβ plaques accumulate in brain areas comprising the DMN due to the high metabolic rate of their continuous activation [59]. This accumulation may disrupt neural mechanisms in structures such as the precuneus and posterior cingulate cortex, which are associated with cognitive processes like memory formation and retrieval; disruptions in these processes may underlie AD symptoms [60].

In our study, we observed associations between cognitive decline and disruptions in two key regions: the hippocampal–entorhinal complex and the precuneus. The hippocampal–entorhinal complex, integral to memory encoding and retrieval, has been consistently implicated in the early stages of AD. Similarly, the precuneus, an integral region of the default mode network, has been associated with metabolic and connectivity changes observed via MRI and [18F]FDG PET in AD [61], further supporting its role in AD-related cognitive decline. Together, these findings suggest that early network-level disruptions in these regions may serve as valuable biomarkers for tracking disease progression and tailoring therapeutic interventions.

The observed alterations in WM networks align with existing literature emphasizing the critical role of WM integrity in cognitive decline associated with the AD course. Previous studies have reported that WM changes may serve as early indicators of neurodegeneration, often preceding GM atrophy [62]. Our research identified significant differences across the WM networks. Specifically, we found notable diagnostic contrasts across cognitive stages, with pronounced changes in WM networks reflecting the underlying pathophysiology of AD. We identified a particularly strong and replicable effect between CN and MCI in the PCoCa. This region has previously been linked to AD, as well as worse cognition [63]. Moreover, PCoCa is on of the structural underpinnings of DMN connection and functioning [64], and it is also impaired at the microstructural level in the different phases of AD continuum [61], demonstrating the ability of high-model-order ICA of amyloid imaging to capture information about the white matter microstructure alterations in AD, further supporting the concept of disconnection mechanisms [65]. The PCoCa plays an important role in sensorimotor functions and many cognitive functions, and lesions in the corpus callosum have been previously associated with cognitive impairment [66]. Our results build upon this and suggest that alterations in WM in the PCoCa may be valuable as a potential biomarker that is prominent in the early stages of AD, as well as for the different clinical presentations of AD-related pathology [67].

With MCI consistently reflecting an intermediate stage between CN and AD, trajectory analysis identified HC-TP and HC-FR as the principal subdomains in which group level loadings for MCI more closely resembled AD than CN, indicating a shift toward an AD-like profile earlier in the disease course. The same directionality was also observed—albeit less consistently—in VI-OT, VI-OC, SC-ET, TN-CE, and TN-DM. This finding suggests that even at the MCI stage, these subdomains may already exhibit advanced or accelerated amyloid burden, potentially foreshadowing more severe neurodegeneration in AD. In contrast, other subdomains placed MCI closer to CN. Collectively, these results highlight a heterogeneity in the vulnerability and progression of amyloid pathology across the brain. Some regions, particularly those crucial for memory encoding and executive function, appear to undergo a more rapid transition toward an AD-like phenotype, whereas others follow a more gradual course. This nuanced view underscores the importance of domain-specific analyses in understanding how different brain networks evolve along the AD course.

Our application of high-model-order ICA enabled us to dissect [18F]FBP PET data into spatially distinct networks, capturing subtle patterns of amyloid deposition across both GM and WM. By leveraging high-order ICA, we moved beyond conventional region-based or voxel-based analyses to reveal fine-grained, network-level disruptions that are not apparent when examining GM or WM in isolation. For instance, while alterations in GM networks, such as those within the hippocampal–entorhinal complex and precuneus, are well-known for their associations with cognitive decline, our approach also revealed distinct WM changes, including in the PCoCa and RICap. Supplementary analyses correcting network loadings for hippocampal volume confirmed that the hippocampal–entorhinal complex within the SC-ET remained significant, and further correction for whole brain volume indicated that AD and CN group differences were largely preserved. Notably, the trajectory of certain MCI subdomains resembling AD rather than CN further underscores ICA’s sensitivity in detecting early transitions, capturing disease progression at a more granular level. This dual-level analysis provides a comprehensive picture of AD pathology, reinforcing the potential of high-order ICA not only to confirm established neurobiological alterations in AD but also to expose novel biomarkers that might lead to more targeted therapeutic interventions.

## Limitations and Future Work

While PET provides valuable molecular insight, it is one of several modalities informative for AD dementia [68]. Our findings align with previous fMRI reports of network alterations [69]. A multimodal fusion strategy that integrates PET (amyloid plus additional tracers such as tau, FDG, translocator protein, etc.) with fMRI [70] and T1-weighted MRI yields complementary molecular, functional, and structural readouts, improving anatomical localization, pathological specificity, and network-level inference. Building on this, we will apply multimodal ICA to fMRI and PET using multi-model-order decompositions to capture effects at multiple spatial scales, considering the granular course of AD (preclinical/prodromal and clinical AD) [71] as well as the different clinical presentation of AD-related pathology (logopenic-variant primary progressive aphasia, posterior cortical atrophy, behavioral variant AD and corticobasal syndrome) [72]. Key limitations as single-tracer emphasis, cross-sectional design, and site/tracer heterogeneity will be addressed via complementary modalities, longitudinal follow-up, and harmonized pipelines, with a focus on predicting conversion from MCI to AD. In addition, CB networks showed decreasing loadings across the trajectory from CN to AD, opposite to the predominantly increasing GM pattern. This decline may reflect an overlap with adjacent WM and smoothing-related partial-volume mixing. The similar decreases observed in WM networks support this interpretation. Future studies should apply higher model orders, cerebellar-specific parcellation, and partial-volume correction to adjudicate CB GM and WM contributions.

In parallel, future work will explicitly examine AD with co-occurring psychosis, which affects a median 41.1% of AD patients (range 12.2–74.1%) and is linked to faster cognitive and functional decline [73]. Psychotic symptoms often overlap with depression (in ∼30–40% of cases), potentially worsening outcomes [74]. These neuropsychiatric features also show notable sex-specific patterns [75], demanding targeted investigation.

## Conclusion

High-model-order ICA of [18F]FBP PET isolates fine-grained amyloid networks in gray and white matter, separating mixed uptake into interpretable components. Differences among CN, MCI, and AD are clearer: HC-TP and HC-FR most consistently placed MCI closer to AD than to CN. Similar patterns were observed intermittently in VI-OT, VI-OC, SC-ET, TN-CE, and TN-DM, while other subdomains showed MCI resembling CN. The hippocampal–entorhinal complex and precuneus have the strongest links to the measures of cognitive decline, consistent with their central roles in AD. The presence of WM PET networks indicates that amyloid-related signal follows distributed pathways rather than being limited to GM. Together, these results show that ICA-based PET network mapping provides anatomically anchored, sensitive measures that sharpen PET-based biomarkers and support disease monitoring.

